# Synthetic hormone-responsive transcription factors can monitor and reprogram plant development

**DOI:** 10.1101/236901

**Authors:** Arjun Khakhar, Alexander R. Leydon, Andrew C. Lemmex, Eric Klavins, Jennifer L. Nemhauser

## Abstract

Developmental programs continuously sculpt plant morphology to meet environmental challenges, and these same programs have been manipulated to increase agricultural productivity^1,2^. Small molecule phytohormones act as signals within these programs creating chemical circuitry^3^ that, in many cases, has been represented in mathematical models^4,5^. To date, model validation and forward engineering of plant morphology has been largely restricted to adding or subtracting genes, as more nuanced tools to modulate key control parameters identified by such models *in vivo* are severely limited^6,7^. Here, we use *Arabidopsis thaliana* to validate a novel set of synthetic and modular <u>h</u>ormone <u>a</u>ctivated <u>C</u>as9-based <u>r</u>epressors (HACRs) that respond to three phytohormones: auxin, gibberellins and jasmonates. We demonstrate that HACRs can regulate genes in response to both exogenous hormone treatments, as well as in response to local differences in endogenous hormone levels associated with developmental events. We further show that HACRs can be used to reprogram the agriculturally relevant traits of shoot branching and phyllotaxy by tuning canalization strength, a critical control parameter predicted by mathematical models. By deploying a HACR to re-parameterize the threshold for induction of the auxin transporter PIN-FORMED1 (PIN1), we observed a decrease in shoot branching and phyllotactic noise as predicted by existing models^4,5^. The approach described here provides a framework for improved mapping of developmental circuitry, as well as a means to better leverage model predictions to engineer development.

## Main text

The body plan of plants is plastic. Extrinsic and intrinsic cues influence developmental programs to allow plants to optimize their form to the environment^3^. Domestication of crops frequently relies on altering these developmental programs to create more productive morphologies for agriculture, such as the dramatic reduction in bushiness of maize^1^ or the dwarfing of cereals that drove the green revolution^2^. Developmental programs are coordinated in large part by a set of chemicals called phytohormones^3^. Accumulation of a given phytohormone by *de novo* synthesis or transport influences the expression or activity of developmental master controllers that go on to direct downstream developmental programs, analogous to wires in a circuit. Auxin, perhaps the best-studied phytohormone, controls many developmental programs that drive agriculturally relevant traits^8^.

Many mathematical models connecting auxin to specific developmental outcomes have been developed^4,5,9^. They highlight the importance of subtle parameters like the strength of feed-forward loops in determining plant morphology. Unfortunately, due to a lack of appropriate tools, these models have only been validated in the extreme states generated by knocking out or overexpressing genes. This has been a major obstacle to leveraging the predictive power of such models to engineer morphologies of agronomic interest. Current plant engineering strategies rely on altering expression of a single gene^7^, an approach ill-suited for tuning the strength of connections within a network.

There is a clear need for tools to re-wire how the phytohormone circuitry regulates developmental programs in plants^10^. However, there are significant hurdles in engineering native chemical circuits. For example, auxin networks are comprised of co-expressed and redundant components, embedded in highly reticulate cross-regulatory relationships with other signaling pathways, and have several layers of feedback^8^, making re-engineering them a daunting challenge. Similar traits are found for most phytohormone-responsive pathways. To facilitate more sophisticated interventions in plant developmental programs, we designed a set of synthetic and modular hormone-activated Cas9-based repressors (HACRs, pronounced ‘hackers’).

We previously validated the design of these synthetic auxin-sensitive transcription factors in *Saccharomyces cerevisiae*^11^. In brief, we fused deactivated Cas9 (dCas9) protein from *Streptococcus pyogenes*^12^ to a highly sensitive auxin-induced degron^13^ and the first 300 amino acids of the TOPLESS repressor (TPL)^14^ (Figure 1A). The dCas9 associates with a guide RNA (gRNA) that targets the HACR to a promoter with sequence complementarity where it can repress transcription. Upon auxin accumulation, the degron sequence targets the HACR for ubiquitination and subsequent proteasomal degradation. Thus, in parallel to the natural auxin response, auxin triggers relief of repression on HACR target genes.

**Figure 1:**
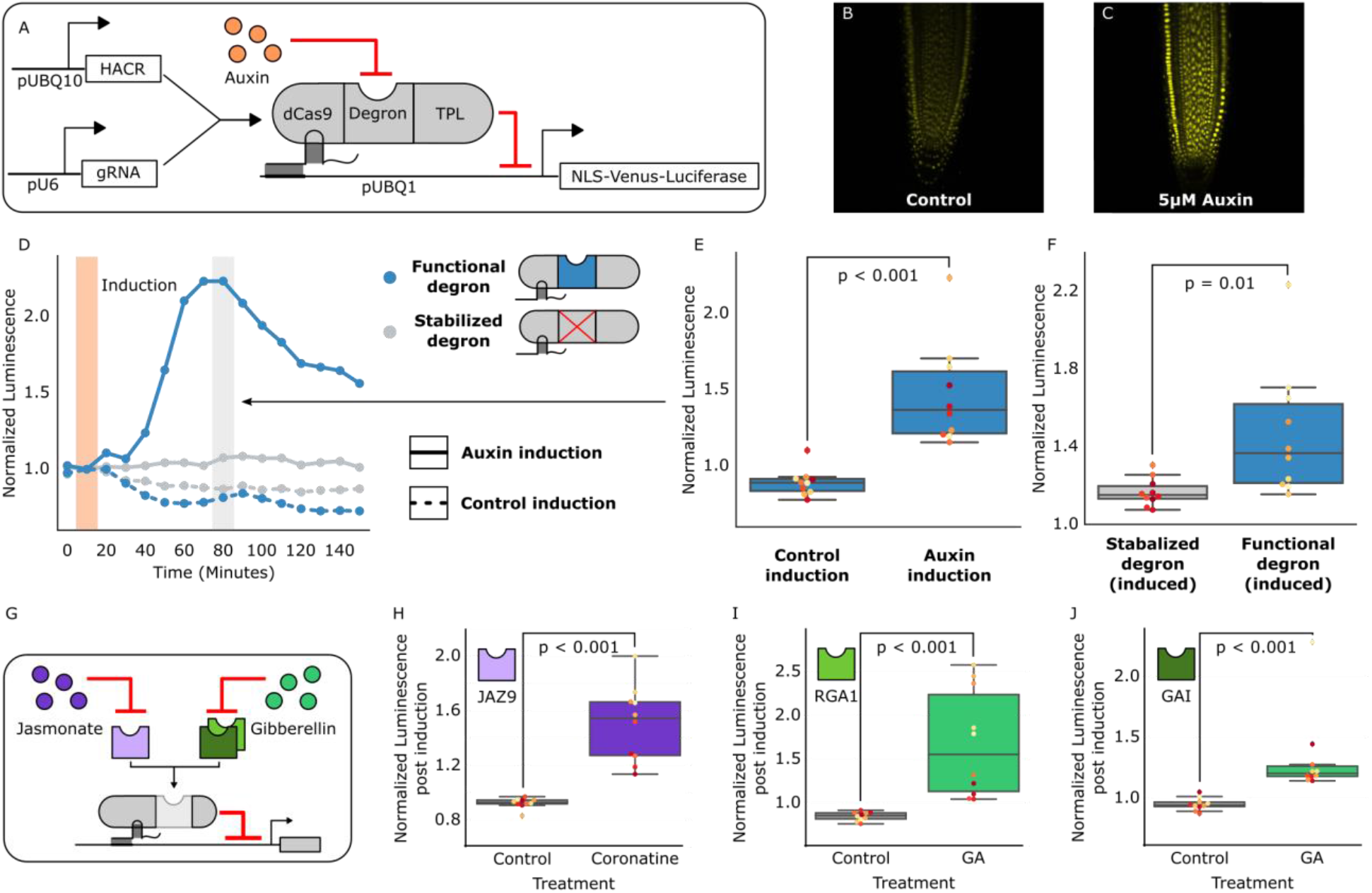
HACRs modulate gene expression upon exogenous hormone treatment. A) A general schematic of the constructs transformed into *Arabidopsis thaliana* to test HACR hormone response. B,C) Confocal microscopy images of root tips from plant lines with an auxin HACR regulating a Venus reporter 24 hours after treatment with (B) control or (C) 5μM auxin. D) An example of a luciferase based time course assay testing whole seedlings of an auxin HACR line treated with auxin (solid blue line) and a control (dashed blue line). The timepoint of auxin induction is highlighted with an orange bar. E) The difference between auxin and control induction at the time of maximum auxin response for the tested seedlings (n = 10) is summarized in the box plot to the right. Every seedling is represented as a different colored dot. F) A HACR variant line with a stabilized auxin degron was also assayed (D, solid and dashed grey lines) and the response to auxin of these seedlings compared to seedlings of the line with a functional auxin degron at the time of maximum auxin response are summarized in box plot in F. G) A schematic of how the hormone specificity of HACRs were altered by swapping the phytohormone degron. H,I,J) These box plots summarize the response of transgenic seedlings carrying these constructs (n=10) to treatment with either control or the appropriate phytohormone. The degron used in the HACR is specified in the top left corner of the plot. Every seedling is represented as a different colored dot. All p-values reported were calculated using a one-way ANOVA.

Transgenic plants were generated with HACRs and a gRNA targeting a constitutively expressed Venus-Luciferase reporter, and, as expected, auxin treatment increased overall fluorescence (Figure 1 B,C). A time-course using luciferase to quantify de-repression of the reporter supported these results with a significant spike in reporter signal (p < 0.001, n = 10) peaking approximately 80 minutes post auxin exposure (Figure 1 D,E). A HACR with a stabilized degron^13^ showed significantly lower reporter signal upon auxin treatment (p = 0.01, n=10) (Figure 1F).

The modular nature of HACRs should allow substitution of the degron with any sequence that has a specific degradation cue. We tested this hypothesis by building HACR variants with degrons sensitive to two other plant hormones: jasmonates (JAs)^15^ and gibberellins (GAs)^16^. Treatment of transgenic plants with exogenous hormones matched to the expressed variants significantly increased reporter signal as compared to control treatments (Figure 1 H, I, J, S1).

To rewire the parameters that connect the phytohormone circuitry with developmental master regulators, HACRs must be able to respond to local differences in endogenous phytohormone levels. To detect subtle differences in HACR sensitivity at the cellular level, we built a ratiometric auxin HACR by combining our previous design with a second reporter (tdTomato) driven by a promoter with a mutation in its gRNA target site (Figure 2A). An estimation of relative auxin levels was then calculated by normalizing the Venus reporter signal in each cell to that of the tdTomato signal in the same cell, minimizing any effect of differential expression of the UBQ1 promoter in different cell types. Using these lines, we visualized tissues at different developmental stages where auxin distributions had been previously described using auxin reporters like DII-VENUS or R2D2^17^. Auxin accumulation assayed by the HACR largely matched previous reports, such as the reverse fountain pattern of reporter signal in the root tip^17^ (Figure 2B) and higher signal in the vasculature as compared to the epidermis of the elongation zone^18^ (Figure 2C). We also observed high reporter signal in emerging lateral root primordia consistent with the auxin accumulation that triggers this developmental event^19^ (Figure D,E).

**Figure 2:**
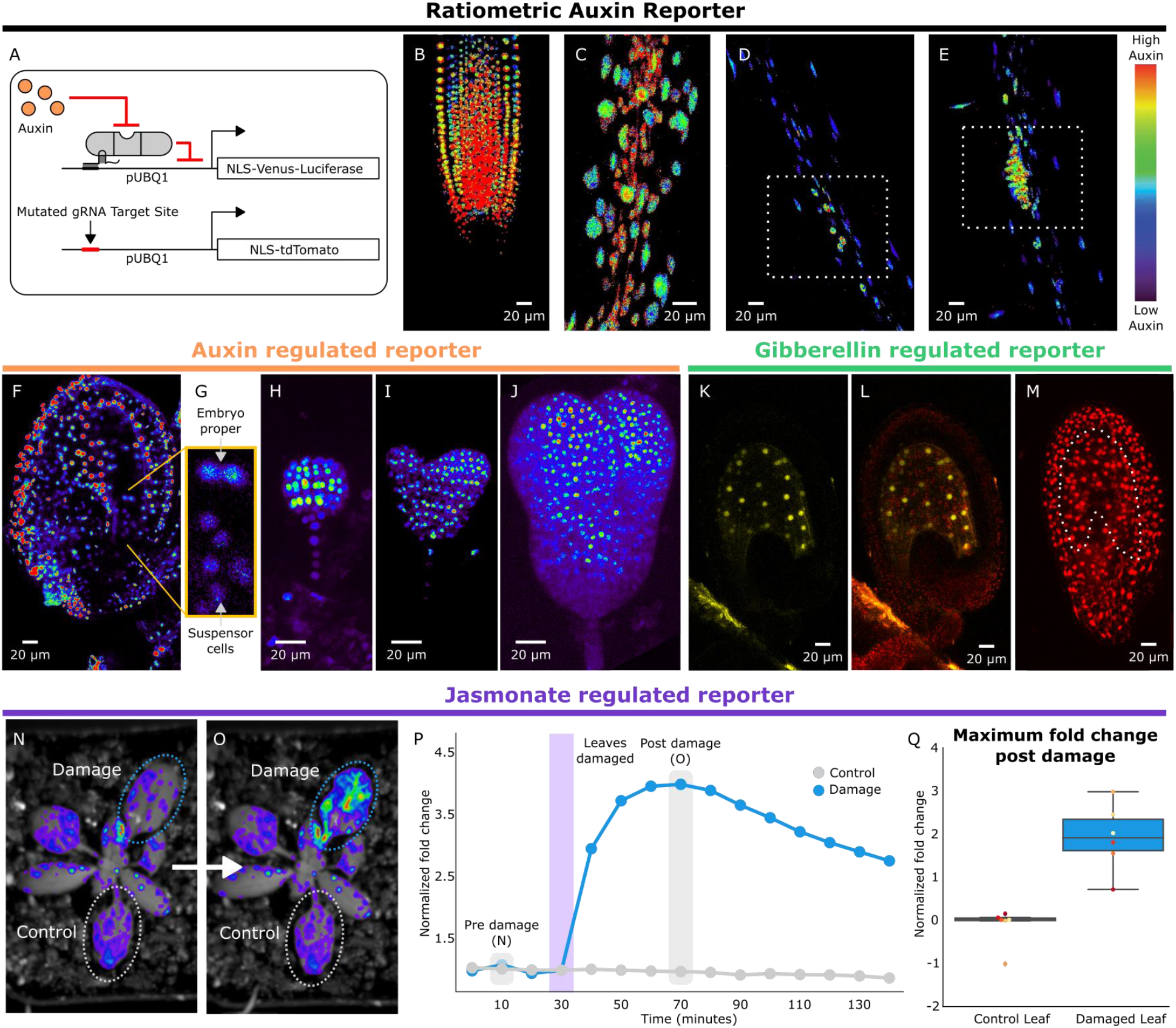
HACRs respond to endogenous hormone signals and can be used to study development. A) Schematic of the genetic circuit used to build ratiometric lines of auxin responsive HACRs. In addition to an auxin HACR regulating a nuclear localized Venus-luciferase reporter the lines also have a nuclear localized tdTomato reporter being driven by a version of the UBQ1 promoter with the gRNA target site mutated. B-E) Confocal microscopy images of roots of seedlings from lines described in A. Reporter signal in images is the background subtracted Venus signal normalized by the background subtracted tdTomato signal. Warmer colors correspond to higher normalized reporter signal. B) The stereotypical reverse fountain pattern of auxin distribution is observed in the root tip. C) Higher reporter signal is observed in the vasculature compared to the epidermis of the elongation zone of the root, consistent with auxin being trafficked along the vasculature. The dashed white boxes highlight high reporter signal in (D) the founder cells of lateral roots and in (E) a developing lateral root primordium. F-J) Confocal microscopy images visualizing reporter signal of a non-ratiometric auxin HACR regulated reporter (F) in the ovule 48 hours post pollination, (G) in the two-cells embryo, (H) in the globular embryo, (I) in the heart stage embryo and (J) in the early torpedo stage embryo. Warmer colors correspond to higher reporter signal. K-M) Confocal microscopy images visualizing reporter signal of a GA HACR regulated reporter (K) in the ovule 48 hours post pollination, (L) reporter signal merged with red auto-fluorescence to highlight the endosperm region and (M) an unregulated tdTomato reporter, with the endosperm highlighted with a dashed white line, for comparison. N-Q) Visualization of JA HACR regulated reporter expression in leaves in response to mechanical damage using a luciferase-based assay. Images of leaves overlaid with the luciferase signal before (N) and after damage (O) are shown to the left of a representative plot of the normalized reporter signal over time (P). Q) Box plot summarizing the maximum fold change at 70 minutes for control and damaged leaves. Points of the same color represent leaves from the same plant.

To further explore the capacity of HACRs to respond to differences in endogenous hormone levels, we visualized the activity of auxin, GA and JA HACRs targeting a Venus reporter. Auxin accumulates in the apical domain of the early embryo and eventually resolves in later stages to the tips of the developing cotyledons, vasculature, and future root apical meristem^17^—the same patterns that were observed in plants expressing an auxin HACR (Figure 2F-J). In plants expressing a GA HACR, we observed strong reporter signal in the early endosperm, consistent with the expression of GA biosynthesis enzymes^20^ (Figure 2KM, S2). There are not many reports of developmental regulation of JA distribution; however, we did detect accumulation of reporter signal in the developing ovule of plants expressing a JA HACR (Figure S2). Specifically, reporter signal appeared to be localized to the inner- and outermost layers of the integuments that surround the developing seed. We also observed that the JA HACR reporter was strongly induced in leaves subjected to mechanical damage (Figure 2N-Q), a condition known to induce high levels of JA^15^. Differences in the patterns of reporter signals in HACR variants demonstrate their specificity for a particular hormone (Figure S2).

Shoot architecture is an agronomically important trait. Fewer side-branches allow for higher density planting^2^ and more regular arrangement of lateral organs (phyllotaxy) facilitates efficient mechanized harvest^21^. The molecular mechanisms that control branching and phyllotaxy are well studied and have been mathematically modeled^4,5^. These models reveal that a key parameter controlling developmental outcome is the strength of canalization, a process by which auxin promotes its own polar transport^22^. The molecular mechanism at the heart of canalization is the auxin-induced increase in levels and activity of the auxin transporter PIN-FORMED1 (PIN1)^22^. To test whether we could alter shoot architecture by decreasing the strength of canalization, we generated transgenic plants with a HACR targeting PIN1 (Figure 3A). In such lines, we would expect that the HACR would increase the threshold level of auxin needed to trigger PIN1 induction, weakening the positive feedback between auxin and PIN1, and thereby reducing canalization strength.

**Figure 3:**
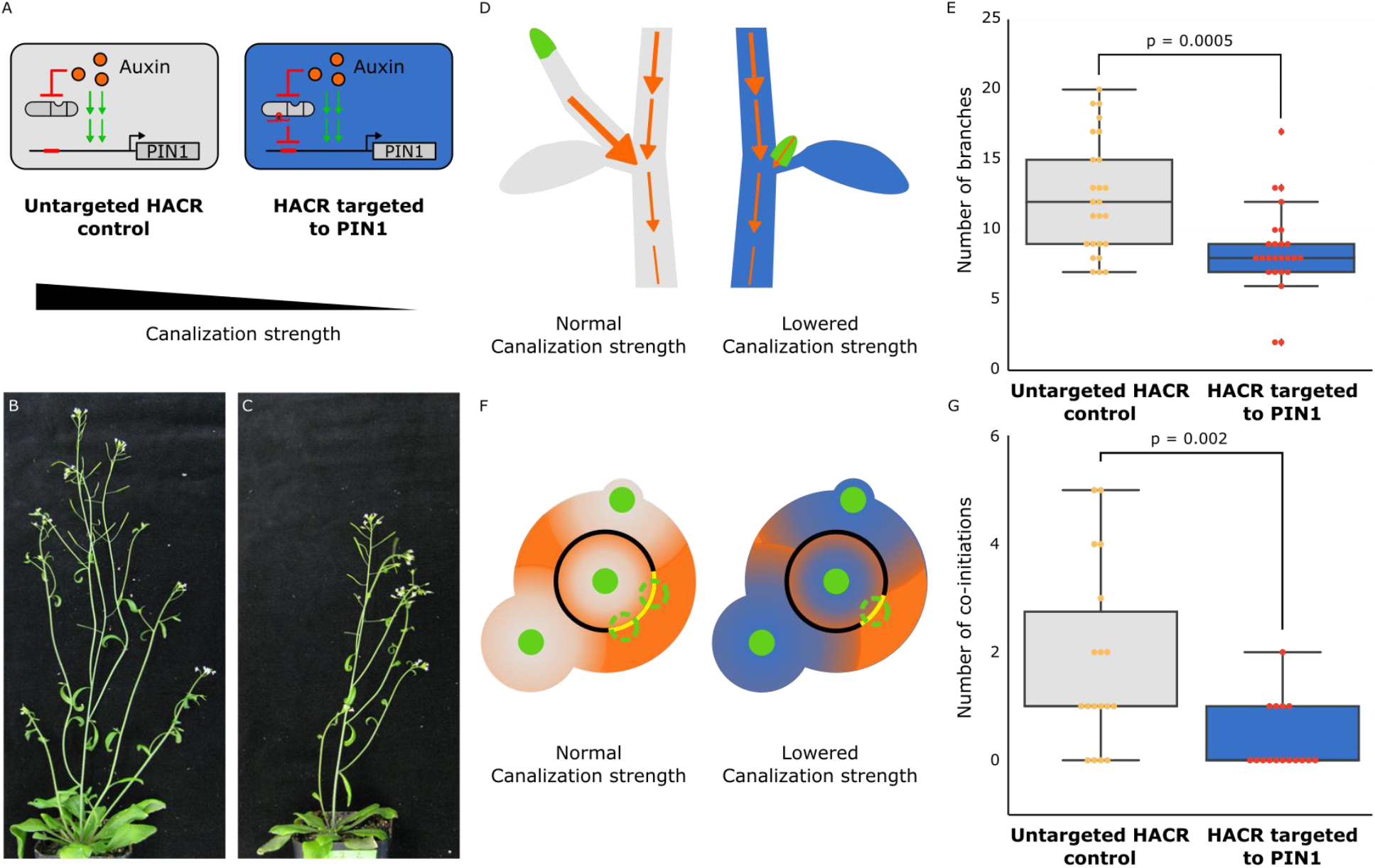
The developmental circuit regulating branching can be rewired using auxin HACRs. A) Schematics of auxin driven PIN1 expression, which is one of the drivers of canalization. In the box on the right we show how we decreased canalization strength by targeting an auxin HACR to regulate PIN1. B,C) Representative pictures of T3 plants of the same age without (B) and with (C) a gRNA targeting an auxin HACR to regulate PIN1. D) Schematic of the mechanism behind the predicted decrease in branching from decreasing canalization strength. In plants without a HACR targeted to PIN1 (grey), the stronger canalization allows the lateral bud (green) to drain auxin (orange arrows) into the central vasculature. In plants with a HACR targeted to PIN1 leading to reduced canalization (blue), the bud is not able to drain its auxin, preventing branch formation. E) Box plots summarizing the number of branches of adult T3 plant lines (n = 25) with a HACR targeted to regulate PIN1 expression (blue boxes), compared to control lines that did not have a gRNA targeting PIN1 (grey boxes). Every dot represents an individual plant. F) Schematic depicting the role of canalization in the pattern of formation of new primordia (green circles) around the shoot apical meristem. In the shoot apex of lines without a HACR targeting PIN1 (grey) the stronger canalization leads to smaller zones of auxin depletion around primordia compared to lines that have a HACR targeting PIN1 (blue). This leads to a broader zone where auxin can accumulate (orange) and create new primordia (dashed green circles) which increases chances of phyllotactic defects. G) Box plots summarizing the number of co-initiations in T3 plant lines (n = 25) with a HACR targeted to regulate PIN1 expression (blue boxes), compared to parental control lines that did not have a gRNA targeting PIN1 (grey boxes). Every dot represents an individual plant. All p-values reported were calculated using a one-way ANOVA.

The model by Prusinkiewicz et al.^5^ simulates how the auxin circuitry regulates branching, and predicts that decreasing canalization strength should result in plants with fewer branches (Supplementary Note 1). This effect is hypothesized to result by the reduced ability of lateral buds to establish auxin efflux into the main stem, an essential step in bud outgrowth (Figure 3D). When we transformed the previously described auxin HACR lines with a construct producing a gRNA targeting the PIN1 promoter, we observed a decrease in the number of branches compared to the parental lines that lacked the gRNA (Figure S3). In lines with the strongest phenotypes, we observed roughly half the total number of branches per plant (Figure 3E). No difference in the number of branches was observed for lines that had a HACR with a stabilized auxin degron regulating PIN1 expression, suggesting this phenotype was not simply due to repression of PIN1 (Figure S4).

Canalization strength is also an important control parameter for the process of phyllotactic patterning. The mechanism driving patterning is hypothesized to rely on each primordium (Figure 3F, green circles) creating a local zone of low auxin that inhibits a new primordia from forming nearby (Figure 3F, shown in orange). This so-called inhibition zone is created by a canalized flow of auxin towards the new primordium. According to a model by Refahi et al.^4^, which captures the stochastic nature of phyllotactic patterning, we would expect that plants with decreased canalization strength would have less noisy phyllotaxy (Supplementary Note 2). The hypothesized mechanism driving this phenotype is that decreased canalization strength should increase the size of the inhibition zone around each primordium, as well as inhibiting the ability of new primordia to establish themselves with effective canalization. Together, these factors would reduce the chances of co-initiating lateral organs. We tested this hypothesis with the plants expressing an auxin HACR targeted to PIN1, and found that indeed these plants had significantly fewer instances of co-initiating lateral organs compared to controls (Figure 3G, S5).

By enabling us to alter the strength of canalization, the HACR platform allowed us to explore previously inaccessible parameter regimes. This proof-of-concept establishes a new method for modifying a large number of desired traits, and provides one of the first demonstrations of model-directed engineering of plant development. The HACR strategy should be extensible to a wide variety of engineered morphologies in both plant and animal systems, particularly given the success of implementing the auxin-induced degradation module (AID) in diverse eukaryotes^23^. In agricultural settings, farmers already manipulate development or defense pathways by applying phytohormones or their synthetic mimics. HACRs could be used to connect these treatments with the expression of genes, such as those involved in defense, to create inducible traits. Additionally, HACRs could be extended to any other phytohormone that utilizes degradation-based signaling, such as salicyclic acid, strigalactones and karrikins. The wide range of degradation cues, the ease of targeting any gene, and the likely conserved function across angiosperms should mean that HACRs have the capacity to reprogram a plethora of developmental traits in a broad range of crop species.

## Methods

### Construction of plasmids

Expression cassettes for the gRNAs, HACRs and the reporters were built using Gibson assembly^24^. These were then linearized by restriction enzyme digestion and assembled into a yeast artificial chromosome based plant transformation vector with kanamycin resistance using homologous recombination based assembly in yeast^25^. The PIN1 gRNA expression vector and the additional tdTomato expression vector for the ratiometric lines were built using Golden-Gate assembly^26^ into the pGRN backbone^27^ with hygromycin resistance.

The gRNA expression cassettes contain a sgRNA driven by the U6 promoter and have a U6 terminator. The HACR expression cassettes are driven by the constitutive UBQ10 (AT4G05320) promoter and have a NOS terminator. All HACR variants contain the same deactivated SpCas9 (dCas9) domain^12^ translationally fused at the N-terminus to an SV40 nuclear localization signal. The phytohormone degron domain and the repressor domain were fused to the C terminus of dCas9, with the respective degron domain in the middle and flexible 6xGS linkers separating the sub-domains. The rapidly degrading NdC truncation of the IAA17 degron^13^ was used for all the auxin HACRs described in the paper. The JA HACR contained the degron from the Arabidopsis JAZ9 protein (AT1G70700)^15^. The GA HACRs contained either GAI (At1g14920)^16^ or RGA1 (At2g01570)^16^ cloned from *Arabidopsis* cDNA. The HACR repression domain was the nucleic acid sequence corresponding to the first 300 amino acids of the TOPLESS repressor (TPL, At1g15750)^14^. We chose this repression domain as TPL is the co-repressor used in native auxin and JA signal transduction pathways. The reporter cassette that was regulated by the HACRs contained a yellow fluorescent protein (Venus) translationally fused to a nuclear localization sequence on its N-terminus and firefly luciferase translationally fused on its C-terminus with flexible linkers. The reporter was driven by a constitutive UBQ1 (AT3G52590) promoter and had a UBQ1 terminator. The additional reporter in the ratiometric lines was identical to these constructs except Venus-Luciferase was replaced with tdTomato and the gRNA target site in the UBQ1 promoter was mutated. The PIN1 gRNA expression vector contained a U6 promoter and terminator.

### Construction of plant lines

All HACR reporter lines were built by transforming the yeast artificial chromosome plasmids described above into *Agrobacterium tumefaciens* (GV3101) and using the resulting strains to transform a Columbia-0 background by floral dip^28^. Transformants were then selected using a light pulse selection^29^. Briefly, this involves exposing the seeds to light for 6 hours after stratification (4°C for 2 days in the dark) followed by a three day dark treatment. Resistant seedlings demonstrate hypocotyl elongation in the case of Hygromycin and leaf greening after 5 days in the case of Kanamycin. After selection seedlings were transplanted to soil and grown in long day conditions at 22°C.

For all the HACR reporter genotypes (Figures 1 and 2) at least three lines were grown to the T2 and tested for their response to the appropriate hormone treatment with n=10 for seedlings. Two different auxin HACR backgrounds were transformed with a gRNA targeting PIN1. The branching of three independent lines, representing three independent PIN1 gRNA insertion events, in each HACR background was characterized in the T2 at n=5. The line with the strongest phenotype was characterized in the T3 at n=25. The number of co-initiations of three independent lines in one HACR background was characterized in the T2 at n=5. The number of co-initiations of one of these lines was characterized in the T3 at n=25.

### Fluorescence Microscopy

For imaging the effects of auxin treatment on root tips we selected plants on 0.5xLS + 0.8% bactoagar containing Kanamycin using the light pulse protocol described above. Four days after the seedlings were removed from the dark we transplanted to fresh 0.5xLS + 0.8% bactoagar without Kanamycin and then imaged on a Leica TCS SP5 II laser scanning confocal microscope on an inverted stand. For auxin induction of root tips, the seedlings were sprayed with a 1:1000 dilution in water of either control (DMSO) or auxin dissolved in DMSO (5μM final concentration) and then mounted on slides in water and imaged after 24 hours.

For the imaging of ratiometric lines seedlings were germinated without selection and then visually screened using a fluorescence microscope for expression of both reporters. These seedlings were then imaged on a confocal microscope at several positions along the primary root to visualize auxin distributions in the root tip, the elongation zone and in developing lateral roots. The images were taken using a Leica TCS SP5 II laser scanning confocal microscope on an inverted stand. The ratiometric images were generated using the calcium imaging calculator in the Leica software, by background subtracting both the tdTomato and Venus signals and then normalizing the Venus signal by the tdTomato signal.

The images of ovules 48 hours after pollination were obtained by emasculating flowers prior to anther dehiscence followed by hand pollination 12 hours after. After 48 hours, the ovules from the pistils of these flowers were dissected using hypodermic needles under a dissection microscope and then mounted on slides in 80mM sorbitol and imaged with confocal microscopy as in Beale *et al*.^30^. To image the developing embryos, ovules were dissected from siliques at the appropriate developmental stages, individually dissected and mounted onto slides in MS0 media before being analyzed by confocal microscopy. All confocal microscopy images presented in this work are maximum projections of sub-stacks from regions of interest.

### Luciferase assays

Luciferase based time course assays were used to characterize the dynamics of HACR response to exogenous or endogenous hormone stimulus. All imaging was done using the NightOWL LB 983 in vivo Imaging System, which uses a CCD camera to visualize bioluminescence. For the data collected for Figure 1 and S1, assays were performed on seedlings. Here, T2 plants were selected by Kanamycin selection using the previously described light pulse protocol. These were then transplanted to fresh plates without antibiotic four days after selection and sprayed with luciferin (5μM in water) in the evening. The next morning, after approximately 16 hours, they were sprayed again with luciferin. After 5 hours they were imaged for one hour (10 minute exposure with continuous time points), then sprayed with a control treatment (a 1:1000 dilution of DMSO in water) and then imaged for five hours. These same plates were then re-sprayed with luciferin (5μM in water) and left overnight. The next day these same plates were again imaged with an identical protocol as the previous day, except they were sprayed with a 1:1000 dilution of hormone in water (5μM Indole-3-acetic acid (auxin), 30μM coronatine (JA) or 100μM GA3 post dilution) rather than control. Luminescence of each seedling was recorded over time and reported as values normalized to the time-point prior to treatment. For the mechanical damage assay of the jasmonate HACR in Figure 2, plants were treated identically as described above except that instead of being sprayed with hormones, leaves on the plant were mechanically crushed using forceps.

### Data Analysis

All the data collected was analyzed and plotted using python (https://github.com/arjunkhakhar/HACR_Data_Analysis). For the luciferase assays, all the time courses were normalized the reading before induction to make them comparable. All p-values reported were calculated in python using the one-way ANOVA function from the SciPy package^31^. (https://docs.scipy.org/doc/scipy/reference/generated/scipy.stats.f_oneway.html)

### Characterizing plant phenotypes

To characterize branching in plant lines with and without an auxin HACR regulating PIN1, we selected T2 transformants for lines that had a gRNA targeting PIN1 and the parental HACR background that had no gRNA. The plants that passed the selection were transplanted onto soil and then characterized as adults at the point that there were on average 4 stems on the no gRNA control lines. In all cases the parental controls that lack a gRNA and the lines derived from them, by transforming with a gRNA targeting PIN1, were all grown in parallel and phenotyped on the same day to ensure the data collected was comparable. Additionally, while we do not believe that the selection would have a significant effect on the phenotyping data as we collected it more than a month after the plants had been transplanted off selection plates onto soil, both the lines with a PIN1 targeting gRNA and the parental controls they were compared to were selected in parallel to control for any confounding effect. Phenotyping involved counting the number of branches on the plant. We quantified the number of branches on five T2 plants for three different lines with a HACR targeted to regulate PIN1 in two different HACR backgrounds, in parallel with the parental HACR background. The line with the strongest phenotype was propagated to the T3 generation with its parental HACR background and the same experiment was repeated with an n=25. To quantify the number of co-initiations we measured the internode length between the first 20 siliques on a single auxiliary stem and every instance of an internode length less than 1 mm was considered a co-initiation. The line that showed the strongest phenotype was propagated to the T3 generation with its parental HACR background and the same experiment was repeated with an n=25.

### Plant genotype list

| Plant genotype | Used in the following figure |
| --- | --- |
| ABS44 (p2301Y-tOCS-pUBQ1:NLS-Venus-LucPlus-tUBQ1-pU6:pUBQ1_gRNA_Target1-tU6-pUBQ10:dCas9-NdC_IAA17-TPLRD2-tNos) | Figure 1B-F , Figure 2F-J , Figure 3B,E,G,H , Supplement Figure 1,2,3,5,6,8 |
| PHD5 (p2301Y-tOCS-pUBQ1:NLS-Venus-LucPlus-tUBQ1-pU6:pUBQ1_gRNA_Target1-tU6-pUBQ10:dCas9-Jas9-TPLRD2-tNos) | Figure 1H, Figure 2N-Q, Supplement Figure 1,2 |
| PHD3 (p2301Y-tOCS-pUBQ1:NLS-Venus-LucPlus-tUBQ1-pU6:pUBQ1_gRNA_Target1-tU6-pUBQ10:dCas9-GAI1-TPLRD2-tNos) | Figure 1J, Figure 2K-M, Supplement Figure 1,2 |
| PHD6 (p2301Y-tOCS-pUBQ1:NLS-Venus-LucPlus-tUBQ1-pU6:pUBQ1_gRNA_Target1-tU6-pUBQ10:dCas9-RGA1-TPLRD2-tNos) | Figure 1I, Supplement Figure 1,2 |
| ABS44 (p2301Y-tOCS-pUBQ1:NLS-Venus-LucPlus-tUBQ1-pU6:pUBQ1_gRNA_Target1-tU6-pUBQ10:dCas9-NdC_IAA17-TPLRD2-tNos)<br>+<br>pGRN_H-pU6:pPIN1_gRNA_Target1-tU6 | Figure 3C,E,G,H, Supplement Figure 3,5 |
| ABS50 (p2301Y-tOCS-pUBQ1:NLS-Venus-LucPlus-tUBQ1-pU6:pUBQ1_gRNA_Target1-tU6-pUBQ10:dCas9-IAA28_DegronDead-TPLRD2-tNos) | Figure 1D,F, Supplement Figure 4 |
| ABS50 (p2301Y-tOCS-pUBQ1:NLS-Venus-LucPlus-tUBQ1-pU6:pUBQ1_gRNA_Target1-tU6-pUBQ10:dCas9-IAA28_DegronDead-TPLRD2-tNos)<br>+<br>pGRN_H-pU6:pPIN1_gRNA_Target1-tU6 | Supplement Figure 4 |

